# The sparse driver system for *in vivo* single-cell labeling and manipulation in *Drosophila*

**DOI:** 10.1101/2024.12.02.626507

**Authors:** Chuanyun Xu, Liqun Luo

## Abstract

In this protocol, we introduce a sparse driver system for cell-type specific single-cell labeling and manipulation in *Drosophila*, enabling complete and simultaneous expression of multiple transgenes in the same cells. The system precisely controls expression probability and sparsity via mutant *FRT* sites with reduced recombination efficiency and tunable FLP levels adjusted by heat-shock durations. We demonstrate that this generalizable toolkit enables tunable sparsity, multi-color staining, single-cell trans-synaptic tracing, single-cell manipulation, and *in vivo* analysis of cell-autonomous gene function.

For details on the use and execution of this protocol, please refer to Xu et al. 2024.

**GRAPHICAL ABSTRACT:** 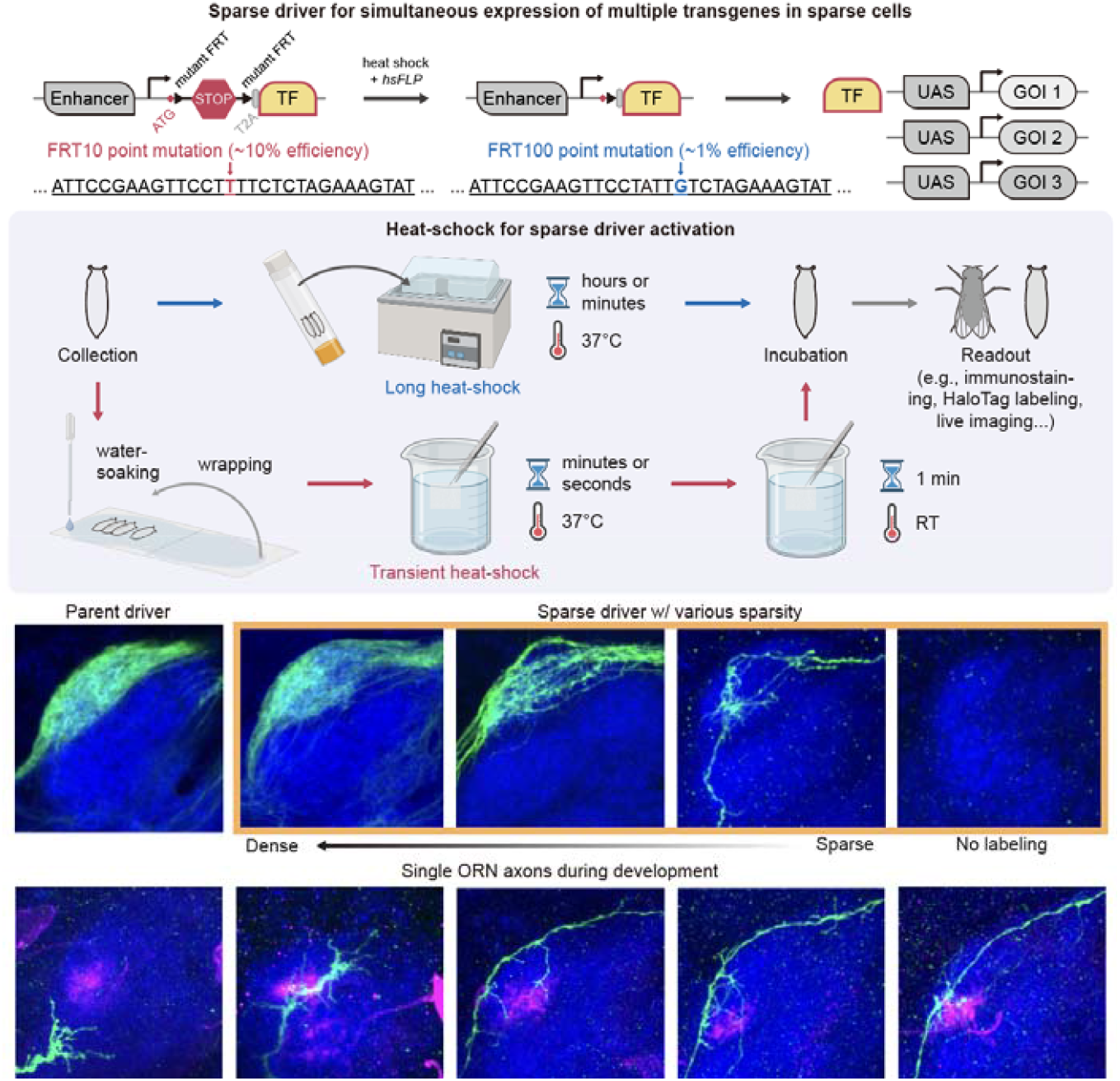

## BEFORE YOU BEGIN

### Background

Sparse neuron labeling and manipulation are powerful tools in neuroscience, allowing for detailed study of individual neurons within complex brain networks (Jefferis and Livet 2012). By targeting a small subset of neurons, researchers can label neuronal morphologies with fluorescent markers, trace synaptic partners with trans-synaptic tracing methods, and monitor real-time activity with GCaMPs. Additionally, by expressing genetic or optogenetic effectors, sparse manipulation enables the study of cell-autonomous gene functions or the dissection of specific neural circuits in behaviors.

However, sparse manipulation methods that rely on probabilistic gating of reporter or effector transgenes often struggle to co-express all desired transgenes in the same subset of neurons (Li et al. 2021; Nern, Pfeiffer, and Rubin 2015; Isaacman-Beck et al. 2020). This issue arises because different reporter or effector transgenes may be activated stochastically in different cell subgroups, as recombination events are independent of each other. A potential solution is to use a stochastically expressed driver transgene that simultaneously controls multiple effectors or reporters, ensuring coordinated expression. The MARCM system (Lee and Luo 1999) is one approach to achieve this. However, MARCM relies on cell division to lose a repressor transgene after mitotic recombination, such that the events cannot be initiated in postmitotic cells. Additionally, its dependence on repressor loss after mitotic recombination hinders its effectiveness in studying developmental events shortly after cell division due to residual repressor activity from mRNA and/or proteins produced before the mitotic recombination event (Wu and Luo 2006).

To address these limitations, we developed a sparse driver system to target single cells within specific neuron types, allowing simultaneous expression of multiple transgenes. Expression probability and desired sparsity are controlled by mutant *FRT* sites with reduced recombination efficiency and tunable FLP recombinase levels through variable heat-shock durations (**Figure 1A**). Point mutations in the *FRT-STOP-FRT* sequence (mutant *FRT10* or *FRT100* sites; Senecoff, Rossmeissl, and Cox 1988) can reduce FLP*-FRT* recombination efficiency by about 10-or 100-fold, respectively (**Figure 1C**). The sparse driver system allows for more precise spatial and/or temporal control, enabling the dissection of cellular events and molecular mechanisms at single-cell resolution.

**Figure 1.**
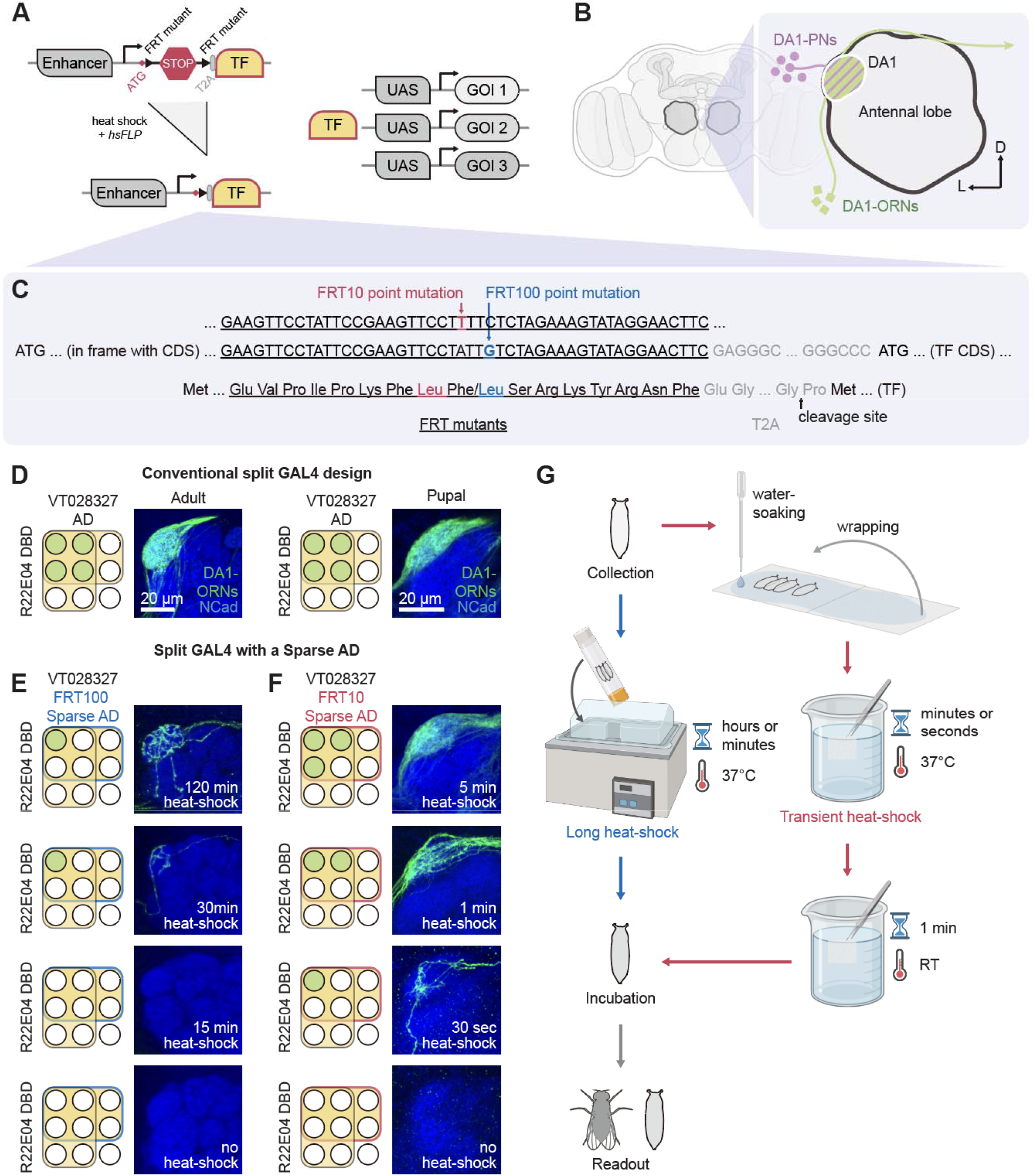
The sparse driver system and its demonstration in the *Drosophila* olfactory circuit. (A) The sparse driver system allows simultaneous expression of multiple transgenes in a subset of cells through stochastic TF (transcription factors) expression. The TF expression is gated by a pair of mutant *FRTs* (*FRT10* or *FRT100* sites) and a transcription termination sequence (shown as *STOP*). Heat-shock-induced stochastic FLP expression removes the *STOP* and enables TF expression in a fraction of cells, driving the co-expression of multiple genes of interest (GOI) in these cells. (B) Adult *Drosophila* brain schematic highlighting antennal lobes and locations of the DA1 glomerulus. Left, DA1-ORN axons (green) synapse with DA1-PN dendrites (purple, contralateral projection omitted). (C) Point mutations (the A→T mutation of *FRT10* or the C→G mutation of *FRT100*) in the *FRT-STOP-FRT* sequence can reduce FLP*-FRT* recombination efficiency by approximately 10-or 100-fold, respectively. Following recombination, the in-frame peptide derived from the mutant *FRT* and *T2A* sequences is excised during the translation of the TF. (D) A conventional split GAL4 strategy to target DA1-ORNs in the adult or pupal antennal lobe. (E) The Sparse^FRT100^-AD-based split GAL4 enables different sparsity tuned by heat-shock time (from 0 to 120 min). (F) The Sparse^FRT10^-AD-based split GAL4 enables different sparsity tuned by heat-shock time (from 0 to 5 min). (G) Two procedures for sparse driver activation.

### *Drosophila* olfactory circuit as a demonstration

In the *Drosophila* olfactory circuit, ∼50 types of olfactory receptor neurons (ORNs) synapse with 50 types of second-order projection neurons (PNs) to form precise 1-to-1 matching at 50 discrete glomeruli (**Figure 1B**), providing an excellent model for investigating mechanisms of synaptic partner matching.

### Driver, reporter, docking site, and mutant *FRT* sequence selection

Effective single-cell morphological characterization requires robust driver and reporter systems. Screening strong drivers and testing reliable reporters (e.g., using *UAS-myr-mGreenLantern* or increasing transgene copies) will improve the signal-to-noise ratio of the following sparse driver experiments. In principle, the sparse driver system works for common driver lines and transcription factors (TFs), e.g., GAL4, QF2, LexA, and their split versions (Luan et al. 2006; Ting et al. 2011; Riabinina et al. 2019). The FlyLight Project (Pfeiffer et al. 2008; Jenett et al. 2012; Tirian and Dickson 2017) has generated extensive anatomical data and well-characterized GAL4, LexA, and Split-GAL4 drivers to visualize and manipulate individual cell types in the *Drosophila* nervous system. If no validated drivers exist for the desired cell type, start with the FlyLight Project database (https://www.janelia.org/project-team/flylight). Notably, since the genomic locations of plasmid docking sites significantly influence driver characteristics (Pfeiffer et al. 2010), select docking sites with expression levels similar to or identical to the original driver for the sparse driver injection. We used the split-GAL4/UAS binary expression system for demonstration, specifically the *VT028327-p65*.*AD* as the parent driver for the sparse driver, along with *GMR22E04-GAL4*.*DBD*, to robustly target DA1-ORN single axons (**Figure 1D**).

***Note:*** The earliest expression timepoint of the sparse driver is controlled by heat-shock timing in experiments and restricted by the original driver’s characteristics. For developmental research, characterize the expression intensities and patterns of the chosen drivers at different developmental stages before designing the sparse driver.

***Note:*** If the properties (e.g., targeted cell number, localization of targeted cells, or expression level) of the parent driver are not well-documented, we recommend testing both *FRT10-STOP-FRT10 (Sparse*^*FRT10*^) and *FRT100-STOP-FRT100 (Sparse*^*FRT100*^) to increase the likelihood of achieving the desired sparsity.

***Note:*** In principle, this protocol can be used in other tissues and different *Drosophila* species. Here, we used the *Drosophila melanogaster* olfactory system for demonstration.

### Experimental model and subject details

Flies (*Drosophila melanogaster*) were raised on standard cornmeal medium in a 12h/12h light cycle at 25°C. For *Sparse*^*FRT10*^, avoid 29°C to prevent any leakiness of *hsFLP*; for *Sparse*^*FRT100*^, 29°C is optional to enhance transgene expression. Details of genotypes used in this study and their sources are described in the **KEY RESOURCES TABLE**.

## KEY RESOURCES TABLE

| REAGENT or RESOURCE | SOURCE | IDENTIFIER |
| --- | --- | --- |
| **Reagents** |  |  |
| Gateway LR Clonase II Enzyme Mix | Thermo Fisher Scientific | Catalog #: 11791020 |
| pENTR/D-TOPO Cloning Kit | Thermo Fisher Scientific | Catalog #: K240020 |
| Zero Blunt TOPO PCR Cloning Kit | Thermo Fisher Scientific | Catalog #: 450245 |
| Phire Tissue Direct PCR Master Mix | Thermo Fisher Scientific | Catalog #: F170L |
| Q5 site-directed mutagenesis kit | New England Biolabs | Catalog #: E0554S |
| Q5 hot-start high-fidelity DNA polymerase | New England Biolabs | Catalog #: M0494S |
| NEBuilder HiFi DNA assembly master mix | New England Biolabs | Catalog #: E2621L |
| **Antibodies** |  |  |
| rat anti-DNcad | Developmental Studies Hybridoma Bank | Catalog #: DN-Ex #8 |
| chicken anti-GFP | Aves Labs | Catalog #: GFP-1020 |
| Rabbit anti-HA | Cell Signaling | Catalog #: 3724S |
| **Chemicals, peptides, and recombinant proteins** |  |  |
| JF646-HaloTag ligand | the Lavis lab | N/A |
| Normal Donkey Serum | Jackson ImmunoResearch | 017-000-121 |
| **Experimental Models: Organisms/Strains** |  |  |
| *D. melanogaster*: GMR22E04-GAL4.DBD | (Jenett et al. 2012) | BDSC: 69199 |
| *D. melanogaster*: VT028327-p65.AD | (Tirian and Dickson 2017) | BDSC: 73064 |
| *D. melanogaster*: QUAS-mtdTomato-3xHA | (Potter et al. 2010) | BDSC: 30004 |
| *D. melanogaster*: trans-Tango | (Talay et al. 2017) | BDSC: 77123 |
| *D. melanogaster*: UAS-mCD8-GFP | (Lee and Luo 1999, 199) | DGRC: 108068 |
| *D. melanogaster*: hsFLP | (Golic and Lindquist 1989) | N/A |
| *D. melanogaster*: UAS-myr-mGreenLantern | (Wong et al. 2023) | N/A |
| *D. melanogaster*: Mz19-QF2<sup>G4HACK</sup> | (Xu et al. 2024) | N/A |
| *D. melanogaster*: UAS-Halo-Moesin | (Xu et al. 2024) | N/A |
| *D. melanogaster*: UAS-V5-Ten-m | (Xu et al. 2024) | N/A |
| *D. melanogaster*: VT028327-FRT10-STOP-FRT10-p65.AD | (Xu et al. 2024) | N/A |
| *D. melanogaster*: VT028327-FRT100-STOP-FRT100-p65.AD | this study | N/A |
| **Recombinant DNA** |  |  |
| pUAS-FRT10-STOP-FRT10-mCD8-GFP | (Li et al. 2021) | N/A |
| pUAS-FRT100-STOP-FRT100-mCD8-GFP | (Li et al. 2021) | N/A |
| pCR-Blunt-TOPO | Thermo Fisher Scientific | Catalog #: K280020 |
| UAS-Halo-CAAX | (Sutcliffe et al. 2017) | Addgene: 87645 |
| VT028327-FRT10-STOP-FRT10-p65.AD construct | (Xu et al. 2024) | N/A |
| VT028327-FRT100-STOP-FRT100-p65.AD construct | this study | N/A |
| **Software and Algorithms** |  |  |
| Zen | Carl Zeiss | RRID: SCR_013672 |
| ImageJ | National Institutes of Health | RRID: SCR_003070 |
| Illustrator | Adobe | RRID: SCR_010279 |

### STEP-BY-STEP METHOD DETAILS

***Note:*** The selection of enhancers, mutant *FRT* sequences, and TFs are highly flexible for preparing the Enhancer-Sparse-TF construct. For simplicity, we use *VT028327-Sparse*^*FRT10*^*-p65*.*AD* as an example.

### Preparation of the Sparse-TF construct

Timing: 1 week

1. Select a plasmid backbone for the desired TF. For example, to generate the DA1-ORN sparse AD, *VT028327-Sparse*^*FRT10*^*-p65*.*AD*, we used *pBPp65ADZpUw* (Pfeiffer et al. 2010).
2. To generate the *FRT10-STOP-FRT10* sequence, PCR amplify *FRT-STOP-FRT* sequence from *pUAST-FRT10-STOP-FRT10-mCD8-GFP*.
3. Insert the *FRT10-STOP-FRT10* sequence and the *T2A* element (to remove the peptide encoded by the *FRT10-T2A* sequence) after the start codon of the p65.AD and keep them in-frame (**Figure 1C**). ***Note:*** Following recombination, the in-frame peptide derived from *FRT10* and *T2A* sequences will be cleaved off from the TF after the translation. ***Alternatives:*** Instead of inserting the *FRT10-STOP-FRT10-T2A* sequence in-frame with the TF coding sequence, an alternative approach is to insert the *FRT10-STOP-FRT10* (without the *T2A* sequence) between the *Drosophila* synthetic core promoter (DSCP) and the coding sequence. Ensure the Kozak sequence remains intact if present in the original construct.
4. Verify the Sparse^FRT10^-AD construct, *pBP-Sparse*^*FRT10*^*-p65ADZpUw*, by sequencing.

### Generation of the Enhancer-Sparse-TF construct

Timing: 1 week

5. Collect primers of selected enhancers from the FlyLight Project. For example, for the enhancer *VT028327*, we first verified its expression in the developing and adult brain and obtained the primer sequences from the project’s website (https://flweb.janelia.org/cgi-bin/view_flew_imagery.cgi?line=VT028327).
6. Lysis the *w1118* strain or the corresponding Bloomington stock (e.g., BDRC #73064 for the VT028327 enhancer) with the Phire Tissue Direct kit (Thermo Fisher).
7. PCR-amplify enhancer fragments from the lysate using primers from the FlyLight Project.
8. Purify the PCR products and confirm their identity through agarose gel analysis and sequencing.
9. Assemble the verified enhancer fragment into the *pENTR/D-TOPO* vector (Thermo Fisher), integrate it into the *pBP-Sparse*^*FRT10*^*-p65ADZpUw* vector using the Gateway LR Clonase II Enzyme Mix (Thermo Fisher), and generate *VT028327-Sparse*^*FRT10*^*-p65*.*AD* construct.
10. Verify the Enhancer-Sparse^FRT10^-AD construct, *VT028327-Sparse*^*FRT10*^*-p65*.*AD*, by full-length plasmid sequencing.

### Generation and maintenance of transgenic flies carrying the sparse driver

Timing: 6–8 weeks

11. Generate transgenic flies in-house using standard methods (Groth et al. 2004) or commercial injection services like BestGene (https://www.thebestgene.com/HomePage.do). Microinject DNA into early *Drosophila* embryos before cellularization. Cross G0 flies to a *white*^*–*^ balancer. Individually balance and verify all *white*^*+*^ progeny. ***Note:*** The Bloomington stock of the parent driver *VT028327-p65*.*AD* uses docking site *attP40*. We selected the docking site *VK00027*, which has a similar expression to *attP40*, for *VT028327-Sparse*^*FRT10*^*-p65*.*AD* and *VT028327-Sparse*^*FRT100*^*-p65*.*AD*. ***Note:*** Keep the sparse driver and *hsFLP* (or other FLP transgenes) in separate stocks. This prevents stochastic FLP expression-mediated recombination and avoids the loss of the *Sparse*^*FRT10*^ or *Sparse*^*FRT100*^ cassette.

### Generation of experimental flies carrying desired transgenes

Timing: 2 weeks

12. Cross parent flies with desired transgenes to get experimental offsprings. ***Note:*** Genotype or functionally validate the sparse driver and other transgenes (e.g., reporters, effectors, *hsFLP*) before the final cross to reduce future troubleshooting difficulties. ***Note:*** The crossing scheme is highly flexible for generating parent flies with the desired transgenes. Adjust the scheme based on the availability and genomic locations of genetic reagents, preferred balancers and markers, and any experimental requirements for a specific gender. ***Optional:*** To assess phenotype in the pupal stage, if possible, use parents with homozygous transgenes or balancers that have markers identifiable in pupae (e.g., *Wee-P* and *Tb*). This is not required but can maximize the likelihood of obtaining the correct genotype in pupae.

### Validation of the sparse driver and titration for desired sparsity

Timing: 1–2 weeks

This procedure requires *Drosophila* samples carrying all necessary transgenes (e.g., reporters, *hsFLP*, sparse driver, with its split partner driver included if required) to target sparse cells.

13. Transfer all adults to a new vial.
14. Remove any existing pupae from the original vial, then set a collection time window to synchronize the pupal stage. ***Note:*** Adjust the collection time window based on the experiment. Longer windows allow more individuals per batch but increase variability in heat-shock-to-readout intervals, leading to inconsistent FLP expression and sparsity. For time-sensitive studies like development, shorter windows are recommended to ensure consistent gene activation timing.
15. After the time window, collect the correct pupae for each test condition (e.g., no heat-shock, 30 min, 1 h, and 2 h of heat-shock) and transfer them into separate vials. ***Note:*** Place them on the wall of each vial and below the water bath water level to maximize the heat transfer efficiency.
16. Heat-shock vials in a 37°C water bath according to the specified durations (**Figure 1G**, left).
17. Wipe up vials and transfer them back to the incubator.
18. Collect pupae or adults at the desired stage. Dissect the flies, perform immunostaining (refer to the **Immunostaining** section), and proceed with imaging.
19. If some heat-shock conditions achieve the desired labeling sparsity, perform an additional round of titration. Determine the optimal heat-shock duration by sampling between the best two conditions, then proceed to the **Sparse labeling and manipulation** section; ***Alternatives:*** Determine the optimal heat-shock duration within current conditions, then proceed to the **Sparse labeling and manipulation** section;
20. If all heat-shock conditions label a large subset of cells, perform another round of fine-tuning titration (e.g., 15 min, 10 min, 5 min, 1 min, or 30 sec durations):
  a. For durations longer than 5 minutes, perform heat-shock in a 37°C water bath as previously described (**Figure 1G**, left; also see steps 15–18). Collect pupae or adults at the desired stage. Dissect the flies, perform immunostaining, and proceed with imaging (**Figure 1E**).
  b. For durations of 5 minutes or less (**Figure 1G**, right):
    1. Wrap pupae in a single layer of paper towel soaked with room-temperature water, ensuring no air bubbles to maintain efficient heat transmission.
    2. Using forceps, immerse the “paper bag” in a 37°C water bath for the target duration, then cool in a room-temperature water bath for 60 seconds.
    3. Transfer the pupae back to the vials and return them to the incubator.
  c. Collect pupae or adults at the desired stage. Dissect the flies, perform immunostaining, and proceed with imaging (**Figure 1F**).
  d. Determine the optimal heat-shock duration within adjusted conditions, then proceed to the **Sparse labeling and manipulation** section.
21. If none of the heat-shock conditions label any cells, refer to the **Troubleshooting** section. ***Note:*** The heat-shock-to-readout interval also affects the accumulated expression level of FLP. Therefore, the heat-shock duration should be adjusted when conducting comparative experiments across early development and adulthood. ***Note:*** The total cell number targeted by the parent driver and the tissue depth will also influence the heat-shock duration required for achieving the desired sparsity. ***Note:*** Since FLP-induced recombination and enhancer activation are independent processes, the heat-shock can be applied before the enhancer activation begins. ***Optional:*** To increase the likelihood and speed of determining the optimal heat-shock duration, test both *Sparse*^*FRT10*^ and *Sparse*^*FRT100*^ in parallel.

### Sparse labeling and manipulation

Timing: 1 week

This procedure requires *Drosophila* samples carrying all necessary transgenes (e.g., effectors, reporters, *hsFLP*, sparse driver, with its split partner driver included if required) to target sparse cells.

***Note:*** In principle, the sparse driver system should allow membrane markers for morphology or live imaging studies, protein markers for subcellular localization studies, tracing tools for trans-synaptic tracing, GCaMPs for real-time activity monitoring, genetic and optogenetic tools for gene and neuron manipulation, or combinations of the above at single-cell resolution.

***Note:*** For demonstration, we used *VT028327-Sparse*^*FRT10/FRT100*^*-p65*.*AD* and *GMR22E04-GAL4*.*DBD* to target DA1-ORN axons with different sparsity (**Figure 1E,F**), *UAS-myr-mGreenLantern* and *UAS-mCD8-GFP* to label the membrane, *Mz19-QF2*^*G4HACK*^ and *QUAS-mtdTomato-3xHA* to orthogonally mark DA1-PN dendrites (**Figure 2A–E**), *trans-*Tango for trans-synaptic tracing (**Figure 2H–H’’**), and *UAS-Halo-Moesin* to label F-actin (**Figure 2I, I’**).

***Note:*** For a protocol outlining how to perform single ORN live imaging, please refer to Li and Luo (2021).

22. Transfer all adults to a new vial.
23. Remove any existing pupae from the original vial, then set a specific collection time window to synchronize the pupal stage. For the DA1-ORN development study, we use a 0–6 h APF (after puparium formation) window.
24. After a 6-hour time window, collect the correct pupae and transfer them into a new vial. ***Note:*** If necessary, select against pupal markers, e.g., *Wee-p* or *Tb*, to maximize the success rate; refer to **Generation of experimental flies carrying desired transgenes** section.
25. For *VT028327-Sparse*^*FRT100*^*-p65*.*AD* (**Figure 1G**, left):
  a. Heat-shock the vial in a 37°C water bath for 1 hour. ***Note:*** Place them on the wall of the vial and below the water bath water level to maximize the heat transfer efficiency.
  b. Wipe up vials and transfer them back to the incubator.
26. For *VT028327-Sparse*^*FRT10*^*-p65*.*AD* (**Figure 1G**, right):
  a. Wrap pupae in a single layer of water-soaked paper towel, ensuring no air bubbles to maintain efficient heat transmission.
  b. Using forceps, immerse the “paper bag” in a 37°C water bath for 30 seconds, then cool in a room-temperature bath for 60 seconds.
  c. Transfer the pupae back to the vials and return them to the incubator.

**Figure 2.**
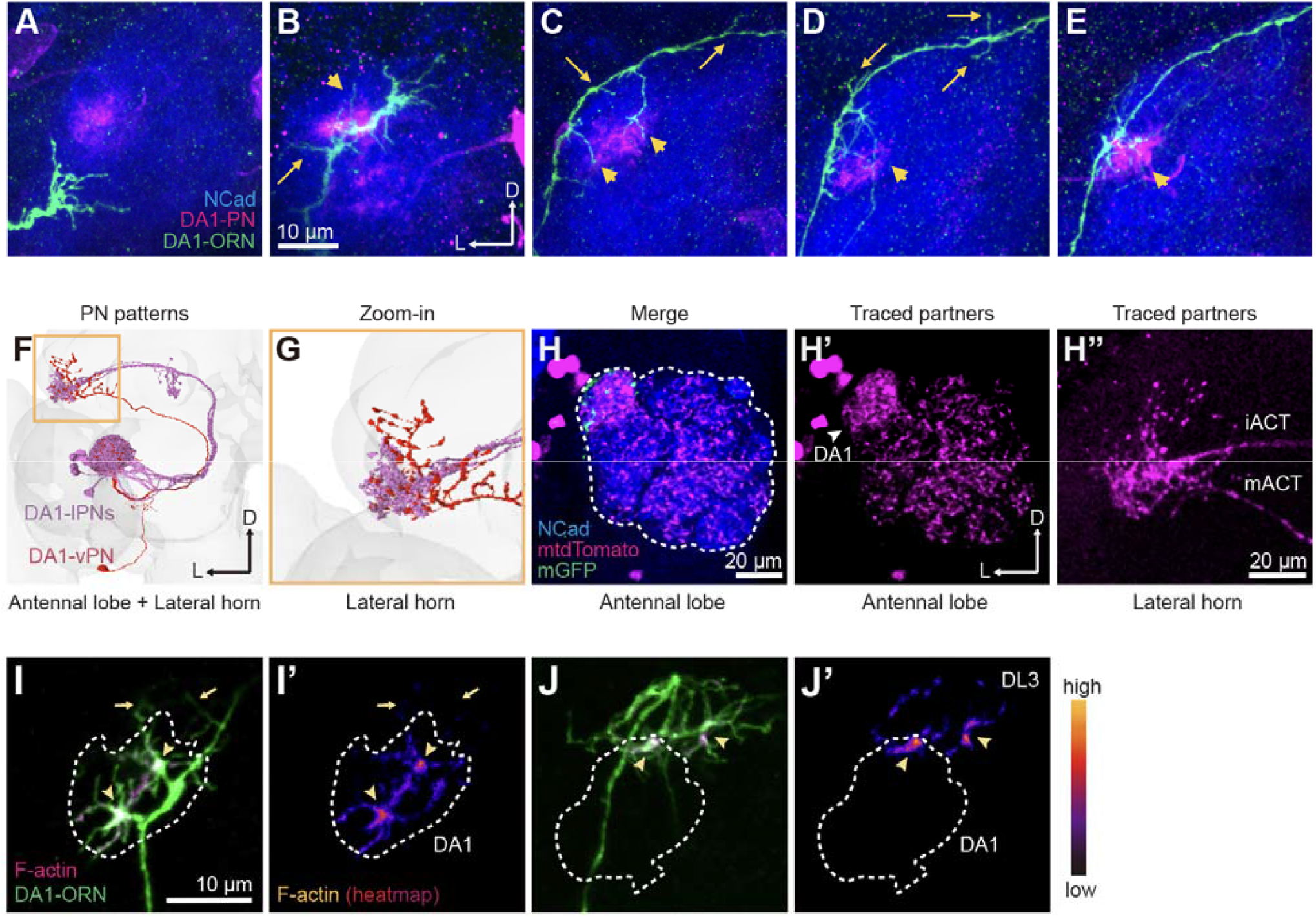
Sparse driver system in various applications. (A–E) Examples of single DA1-ORN axons (green) innervating the ipsilateral antennal lobe in different stages. Axonal exuberant branches that contact target DA1-PN dendrites (magenta) are eventually stabilized, showing a stabilization-upon-contact manner. Arrows, non-DA1-PN-contacting (OFF-target) branches; arrowheads, DA1-PN-contacting (ON-target) branches. (F, G) FlyWire tracings of DA1-lPNs and DA1-vPN from the left hemisphere (F), with a magnified view at the lateral horn (yellow box), to visualize their stereotyped axon branching patterns (G). (H-H’’) Representative confocal images of *trans-Tango*-mediated trans-synaptic tracing from DA1-PNs. Green, ORN axons; magenta, postsynaptic neurons labeled by *trans-Tango*, which include dendrites of local interneurons and more intensely labeled DA1-PNs in the antennal lobe (H, H’) and DA1-PN axons in the lateral horn (H’’). Dashed outlines, antennal lobe. (I, I’) Representative confocal images of the F-actin distribution (I, magenta; I’, heatmap based on Halo-Moesin staining) in a control DA1-ORN axon (I, green). Arrows, non-DA1-PN-contacting subregion; arrowheads, F-actin hotspots. Dashed white traces outline DA1-PN dendrites. (J, J’) Representative confocal images of the F-actin distribution in a Ten-m overexpressing DA1-ORN axon (targeting to the DL3 glomerulus). Labels are the same as I and I’. D, dorsal; L, lateral. NCad (N-cadherin) is a general neuropil marker.

### Immunostaining

Timing: 5 days

27. Collect pupae or adults at the desired stage.
28. Dissect brains or other tissues in pre-cooled phosphate-buffered saline (PBS).
29. Fix them in 4% paraformaldehyde in PBS with 0.015% Triton X-100 for 15 minutes on a nutator at room temperature. ***Note:*** If necessary, adjust fixation conditions to minimize background from over-fixation.
30. Wash the fixed brains with PBST (0.3% Triton X-100 in PBS) four times, nutating for 15 minutes each time.
31. Block the brains in 5% normal donkey serum in PBST (blocking solution) for 1 hour at room temperature or overnight at 4°C on a nutator.
32. Dilute primary antibodies in the blocking solution and incubate the brains with the antibodies for 36–48 hours on a 4°C nutator. ***Note:*** Primary antibodies used in immunostaining include rat anti-NCad (1:40; DN-Ex#8, Developmental Studies Hybridoma Bank), chicken anti-GFP (1:1000; GFP-1020, Aves Labs), and rabbit anti-HA (1:100, 3724S, Cell Signaling).
33. After incubation, wash the brains with PBST four times, nutating for 20 minutes each time.
34. Incubate the brains with secondary antibodies diluted in the blocking solution, nutating in the dark for 24–48 hours at 4°C. ***Note:*** Donkey secondary antibodies conjugated to Alexa Fluor 405/488/568/647 (Jackson ImmunoResearch or Thermo Fisher) were used at 1:250.
35. Rewash the brains with PBST four times, nutating for 20 minutes each time.
36. Mount the immunostained brains with SlowFade antifade reagent and store them at 4°C until imaging.

### HaloTag labeling (optional)

Timing: 1 day

37. Dissect fly brains in pre-cooled PBS and fix them in 4% paraformaldehyde in PBS for 10 minutes on a nutator at room temperature.
38. Wash the fixed brains with PBST for 5 minutes, repeating the wash thrice. Incubate the brains with Janelia Fluor 646 HaloTag Ligand (0.5 μM in PBS) for 5 hours or overnight at room temperature in the dark.
39. After incubation, wash the brains with PBST for 5 minutes, repeating three times.
40. If needed, proceed with the immunostaining protocol (refer to the **Immunostaining** section, steps 31–36).

## EXPECTED OUTCOMES

The example dataset includes sparse axons of DA1-ORNs in the developing or adult antennal lobe.

### Axon labeling with different sparsity in the adult or pupal brain

Across a range of heat shock durations (15 to 120 minutes), DA1-ORN Sparse^FRT100^-AD-based split GAL4 labeled no axons or sparse axons in the adult antennal lobe (**Figure 1E**). Over 0.5–5 minutes of heat shock, Sparse^FRT10^-AD-based split GAL4 labeled a single axon, an intermediate subset, or a large subset of DA1-ORN axons (**Figure 1F**).

### ORN-PN synaptic partner matching at single-axon resolution

The orthogonal labeling of the DA1 ORN-PN pair revealed that, during development, the DA1-ORN axon initially overproduces exuberant branches along the stem axon to expand the searching space for target selection (**Figure 2A–E**). Over time, branches that contact the target PN dendrites are stabilized, while OFF-target branches are pruned.

### Single neuron trans-synaptic tracing

*Trans-*synaptic tracing of a single DA1-ORN axon labeled its contacting neurons, including DA1-PNs and local interneurons, in the antennal lobe (**Figure 2H-H’’**). The traced DA1-PNs exhibited a morphology similar to DA1-PNs reconstructed in the Flywire database (**Figure 2F, G**).

### Single axon manipulation and HaloTag staining

Co-labeling of membrane marker and F-actin marker in control DA1-ORN single axons revealed that contact with DA1-PN dendrites promoted local F-actin levels in target-contacting branches (**Figure 2I, I’**). Overexpression of Ten-m (tenascin-major) (Hong, Mosca, and Luo 2012; Xu et al. 2024), a transmembrane protein instructing synaptic partner matching, in single axons led to axon mistargeting and promoted F-actin levels in the DL3 glomerulus (**Figure 2J, J’**).

***Note:*** Data from **Figure 1D, F**, and **Figure 2** are reprinted with permission from Xu et al. 2024.

| Fly genotype |  |
| --- | --- |
| Figure 1D | *UAS-dcr2, UAS-CD8-GFP / +; VT028327-p65.AD / +; GMR22E04-GAL4.DBD / +* |
| Figure 1E | *UAS-CD8-GFP, hsFLP / UAS-dcr2, UAS-CD8-GFP; ; GMR22E04-GAL4.DBD / VT028327-FRT100-STOP-FRT100-p65.AD* |
| Figure 1F | *UAS-CD8-GFP, hsFLP / UAS-dcr2, UAS-CD8-GFP; ; GMR22E04-GAL4.DBD / VT028327-FRT10-STOP-FRT10-p65.AD* |
| Figure 2A–E | *UAS-CD8-GFP, hsFLP / UAS-dcr2, UAS-CD8-GFP; Mz19-QF2<sup>G4HACK</sup>, QUAS-mtdTomato-3xHA / UAS-myr-mGreenLantern; GMR22E04-GAL4.DBD / VT028327FRT10-STOP-FRT10-p65.AD* |
| Figure 2H–H'' | *QUAS-mtdTomato-3xHA, UAS-CD8-GFP, hsFLP / + ; trans-TANGO / +; GMR22E04-GAL4.DBD / VT028327-FRT10-STOP-FRT10-p65.AD* |
| Figure 2I, I' | *UAS-CD8-GFP, hsFLP / UAS-dcr2, UAS-CD8-GFP; Mz19-QF2G4HACK, QUAS-mtdTomato-3xHA / UAS-myr-mGreenLantern; GMR22E04-GAL4.DBD / VT028327-FRT10-STOP-FRT10-p65.AD, UAS-Halo-Moesin* |
| Figure 2J, J' | *UAS-CD8-GFP, hsFLP / UAS-dcr2, UAS-CD8-GFP; Mz19-QF2G4HACK, QUAS-mtdTomato-3xHA / UAS-myr-mGreenLantern; GMR22E04-GAL4.DBD, UAS-V5-Ten-m / VT028327-FRT10-STOP-FRT10-p65.AD, UAS-Halo-Moesin* |

## LIMITATIONS

Although this protocol enables single-cell visualization and manipulation *in vivo*, it has a few limitations. First, the performance of the sparse driver system relies heavily on the properties of the parent driver. If the parent driver fails to target the desired cell population effectively, this protocol cannot robustly access single cells within that population *de novo*. Additionally, the sparse driver system has only been tested with enhancer lines from the FlyLight Project; GAL4-based enhancer trap lines and T2A-GAL4 knock-in lines remain untested. Second, while heat-shock at the pupal stage ensures controllable heat transmission, heat-shock at the larval or adult stage has not been tested. Third, the heat-shock promotor may respond to other stimuli, such as heavy metals, oxidative stress, UV radiation, hypoxia, inflammation, and certain chemical treatments. Therefore, this protocol may not be suitable for experiments involving these treatments.

## TROUBLESHOOTING

### Problem 1

No cells are labeled after sparsity titration.

### Potential solution

If a 2-hour heat-shock duration does not label any cells:

a. Try extending the duration or performing multiple heat-shocks.
b. Verify the genotype of experimental pupae, parents, and stocks to ensure all necessary components are present.
c. Sequence the sparse driver stocks to confirm the presence of both mutations in either *FRT10-STOP-FRT10* or *FRT100-STOP-FRT100*.
d. Functionally validate *hsFLP* by using *UAS-FRT-STOP-FRT-mCD8-GFP*. Double-check the characteristics of the original driver to confirm if it is active at the assay stage.
e. Use stronger parent drivers (if available), improved reporters (e.g., *UAS-myr-mGreenLantern* and *UAS-Halo-CAAX*), and optimized fixation and staining conditions to enhance the signal-to-noise ratio.
f. Increase the sample size of each condition.

### Problem 2

Too many cells are labeled after sparsity titration.

### Potential solution

If a 30-second heat-shock duration still labels too many cells:

a. Try reducing the duration or decreasing the heat-shock temperature.
b. Try the *FRT100-STOP-FRT100* sequence.
c. Reduce the interval between the heat-shock and the readout time.
d. Sequence the sparse driver stocks to ensure that the *FRT10-STOP-FRT10* or *FRT100-STOP-FRT100* have been correctly constructed.

### Problem 3

Samples show a high background or weak signal.

### Potential solution

Use stronger parent drivers (if available), improved reporters (e.g., *UAS-myr-mGreenLantern* and *UAS-Halo-CAAX)*, and optimized fixation and staining conditions to enhance the signal-to-noise ratio.

### Problem 4

The probability of the same sparsity is inconsistent across batches.

### Potential solution

a. Ensure that intervals between the heat-shock and the readout time are consistent across batches.
b. Confirm that transgenes are homozygous in the parent generation or strictly select against pupal markers, e.g., *Wee-p* or *Tb*, in the experimental generation.
c. Perform consistent heat-shock durations across batches.
d. If the heat-shock duration is 5 minutes or less, ensure no air bubbles in the “paper bag” and sufficient cooling time in the room temperature water bath.
e. Check the copy number of *hsFLP* transgene.

### Problem 5

Animals are dying after long heat-shock.

### Potential solution

Weaker flies may need multiple shorter heat-shocks; try 2–3 heat-shocks of 30 minutes each, with 30-minute recovery intervals.

### Problem 6

Labeled cells in the control condition without heat-shock.

### Potential solution

Normally, in the absence of FLP, the sparse driver does not spontaneously recombine. Spontaneous recombination is typically caused by FLP leakage and accumulated expression. To address this, reduce the heat-shock-to-readout interval for all conditions (ensuring consistency in experimental timing), use the less sensitive *FRT100-STOP-FRT100*, or raise flies at 25°C.

### Problem 7

The expression level of a transgene changes when co-expressed with varying numbers of other transgenes or between experimental and control groups.

### Potential solution

When a cell expresses multiple transgenes, they share the same driver TFs. If the driver cannot produce enough TFs for all transgenes, the TF dilution effect becomes apparent. To ensure consistency, maintain the same number of transgenes across conditions. Ideally, prepare a control transgene inserted into the same locus as the key transgene whose biological effect is to be examined.

## RESOURCE AVAILABILITY

### Lead contact

Further information and requests for resources and reagents should be directed to and will be fulfilled by the lead contact, Chuanyun Xu.

### Materials availability

Plasmids and *Drosophila* lines are available upon request from the lead contact.

### Data and code availability

This study did not generate or analyze datasets or code.

## ACKNOWLEDGMENTS

We thank members of the Luo lab, especially Z. Li, C. Lyu, D.C. Wang, Y. Zhang, and C. McLaughlin, for support, insights, and feedback on this project. Figures were partially created using BioRender.com. Supported by an NIH grant (R01-DC005982 to L.L.).

## AUTHOR CONTRIBUTIONS

Methodology and Investigation, C.X.; Writing, C.X. and L.L.; Funding Acquisition and Supervision: L.L.

